# The impact of mutations on the structural and functional properties of SARS-CoV-2 proteins: A comprehensive bioinformatics analysis

**DOI:** 10.1101/2021.03.01.433340

**Authors:** Aqsa Ikram, Anam Naz, Faryal Mehwish Awan, Bisma Rauff, Ayesha Obaid, Mohamad S. Hakim, Arif Malik

**Affiliations:** Institute of Molecular Biology and Biotechnology (IMBB), The University of Lahore (UOL), Lahore, Pakistan; Department of Medical Lab Technology, University of Haripur (UOH), Haripur, Pakistan; Department of Microbiology, University of Haripur (UOH), Haripur, Pakistan; Department of Microbiology, Faculty of Medicine, Public Health and Nursing, Universitas Gadjah Mada, Yogyakarta, Indonesia; Center for Child Health – PRO, Faculty of Medicine, Public Health and Nursing, Universitas Gadjah Mada, Yogyakarta, Indonesia

**Keywords:** mutation, RdRp, spike, SARS-CoV-2, 3CLpro

## Abstract

An in-depth analysis of first wave SARS-CoV-2 genome is required to identify various mutations that significantly affect viral fitness. In the present study, we have performed comprehensive in-silico mutational analysis of 3C-like protease (3CLpro), RNA dependent RNA polymerase (RdRp), and spike (S) proteins with the aim of gaining important insights into first wave virus mutations and their functional and structural impact on SARS-CoV-2 proteins. Our integrated analysis gathered 3465 SARS-CoV-2 sequences and identified 92 mutations in S, 37 in RdRp, and 11 in 3CLpro regions. The impact of those mutations was also investigated using various in silico approaches. Among these 32 mutations in S, 15 in RdRp, and 3 in 3CLpro proteins are found to be deleterious in nature and could alter the structural and functional behavior of the encoded proteins. D614G mutation in spike and P323L in RdRp are the globally dominant variants with a high frequency. Most of them have also been found in the binding moiety of the viral proteins which determine their critical involvement in the host-pathogen interactions and drug targets. The findings of the current study may facilitate better understanding of COVID-19 diagnostics, vaccines, and therapeutics.

## Introduction

Severe acute respiratory syndrome coronavirus 2 (SARS-CoV-2), among the seventh known human infecting coronaviruses, is a highly transmissible and pathogenic virus [1]. It belongs to the *Betacoronavirus* genus and is enveloped, positive-sense, and single-stranded RNA virus [2]. Mutation is a distinct aspect of RNA viruses depending upon the fidelity of their RNA-dependent RNA polymerase (RdRp) [3]. Mutation can have their advantages for viruses and can contribute to viral adaptation towards pathogenesis, immune escape, and drug resistance [4]. Many mutations cause drug resistance and affect the virulence of various pathogenic viruses. Hence, they have a great impact on human health, speculating that any new mutations in SARS-CoV-2 can be hazardous during this rapidly escalating outbreak. Studies performed over the past few months have revealed and suggested that SARS-CoV-2 have some evolving mutations in their human host [5].

The functional and structural consequences of these mutations are unknown, and it will be substantial to determine its impact on transmissibility and pathogenesis in humans. The analysis of genetic sequence data freely available at NCBI (https://www.ncbi.nlm.nih.gov/nuccore) and Global Initiative on Sharing All Influenza Data (GISAID; https://www.epicov.org) sheds light on key epidemiological parameters of SARS-CoV-2, including evolving mutations globally [6]. Therefore, we kept our focus on SARS-CoV-2 mutations lying within RdRp, 3C-like protease (3CLpro), and spike proteins in an attempt to assess the spread of new viral variants across the countries and also the real functional and structural impact of these mutations on the pathogenicity of SARS-CoV-2. These viral proteins are considered among the primary targets for vaccine and antiviral drug development.

A more comprehensive understanding of virulence mutations and their evolution can be achieved by genomic analysis of sequence data that can further guide to various experimental studies. The availability of such comprehensive data is enabling researchers to use various bioinformatics tools in an attempt to extract useful hidden clinical and molecular information [7]. However, there is a need to uncover deleterious mutations and their pathogenic variants from the readily available data and to further explore their impact at the molecular level. *In-silico* tools can be effectively utilized for prioritizing different variations in a cost-efficient manner and to further investigate structural and functional consequences of specific mutations [8]. Thus, in this study all available genomic information of the first wave of SARS-CoV-2 outbreak have been retrieved and various *in-silico* approaches have been used to provide an insight into the pathogenic landscape of various mutations on selected viral proteins.

The main aim of the study was to understand and predict various pathogenic variants in the first wave of SARS-CoV-2 RdRp, 3C-like protease (3CLpro), and spike (S) proteins. Overall, 32 mutations in S, 15 in RdRp, and 3 in 3CLpro have been predicted in this study, which are involved in major phenotypic damage and could alter the structural and functional behavior of the encoded proteins.

## Material and Methods

### 1. Sequence retrieval

All complete genome sequences of the first wave SARS-CoV-2 (n=3465) were downloaded from the GenBank and GISAID until 15^th^July, 2020. Genome sequence (NC_045512) was used as a reference sequence and is considered as a wild type (WT) sequence. From these complete genome sequences, sequences of S, RdRp and 3CLpro regions were screened out.

### 2. Sequence alignment and mutation analysis

Protein sequences of S, RdRp and 3CLpro regions were first aligned with the reference sequence (NC_045512) using CLC workbench 7 and Bioedit [9]. The origin and position of each mutation within these viral proteins are assessed.

### 3. The impact of mutation on the structure and functional properties of the encoded viral proteins

Prediction of different mutations that alters the structure and functions of SARS-CoV-2 proteins can actually guide designing pharmaceutical compounds and initiate the vaccine design and development. Thus, to estimate the effect of the identified mutations on various structural and functional features of COVID-19 viral proteins, following analyses were performed:

### 3.1. Predicting the functional impact of mutations

To characterize mutations as neutral or deleterious to the structure and function of the encoded proteins, SIFT [10], PhD-SNP [11], and SNAP2 tools [12] were employed. SIFT predicts the functional importance of an amino acid variations based on the conservation and alignment of highly similar orthologoue and paralogoue protein sequences. Substitutions with probability score less than 0.05 are considered to be deleterious, while values ≥0.05 are considered to be tolerated, i.e. they may have no significant effect.

PhD-SNP is a support vector machine-based software and predict whether the substitution may cause a disease or may remain neutral. The SNAP2 (screening for non-acceptable polymorphisms) program (www.rostlab.org/services/SNAP/) makes predictions regarding the functionality of variant proteins.

### 3.2. Predicting protein stability changes upon mutations

Prediction of the mutations impact on the conformation, flexibility, and stability of protein is also required to gain an insight into structure-function relationship of the encoded protein. Protein stability is the basic characteristic that affects the function, activity, and regulation of proteins [13]. Free energy related to protein unfolding is a key index of protein stability. Therefore, by analyzing the influence of mutation on free energy, its effect on protein stability could be accurately determined. To quantitatively predict the change in protein conformation, flexibility, and stability due to mutations, i-Mutant version 2.0 [11], DUET [14] and Dynamut [13] web servers were used. For DUET and Dynamut prediction, 3D structure of RdRp and S were predicted using i-TASSER while crystal structure (5re5) of 3CLpro was retrieved from protein data bank (PDB).

## 4. Mutation screening

In order to recapitulate the predictive results of above-mentioned tools, a scoring criterion was set (0-6). If a mutation were predicted to be “harmless” or “neutral” by all tools, it would score 0. Though, it would get a score if any of the tool predicted it as a “harmful” or “Pathogenic” mutation respective of the number of tools predicting it. Mutations predicted by four or more tools (thus with score ≥4) were then screened for further evaluations.

## 5. Normal mode analysis

Each deleterious mutation (score ≥4) was inserted in the PDB structure of each selected viral protein by using chimera and Normal mode analysis was performed via iMod server (iMODS) (http://imods.chaconlab.org) by using the basic default values for all the parameters mentioned.

## 6. Mapping the ligand binding sites with mutations

To find the location of screened mutations within the drug binding sites of viral proteins, COACH (http://zhanglab.umich.edu/COACH/) and CASTP (http://sts.bioe.uic.edu/castp/index.html?2r7g) servers were used. These servers predict protein-ligand binding sites and thus these sites were evaluated for the presence of any pathogenic mutations. Mutations lying within these regions were then screened out to have negative effect on the targeted proteins and their possible interactions.

## Results

### Mutations residing in S, RdRp, and 3CLpro sequences

Alignment of 3,465 SARS-CoV-2 protein sequences with the reference sequence Wuhan-Hu-1 (Accession NC_045512) revealed 92 mutations in S, 37 in RdRp, and 11 in 3CLpro regions (Table 1 and Figure 1). These mutations were found to be in a wide range of countries, including the United States (US), China, Australia, South Korea, India, Peru, Sweden, Spain, Vietnam, England, Pakistan, Turkey, Germany, France, Greece, Sri Lanka, South Africa, Colombia, Iran, and Malaysia. It indicates that the virus has a significantly high evolution rate in various geographical regions to increase the viral fitness. D614G (50%) and P323L (49%) mutations showed the highest frequency among the screened sequences (n=3,465). Moreover, the mutation frequencies of P323L (49%) and D614G (50%) was found to be similar within the period of 15 January 2020 to 15^th^ July 2020.

**Table 1.**
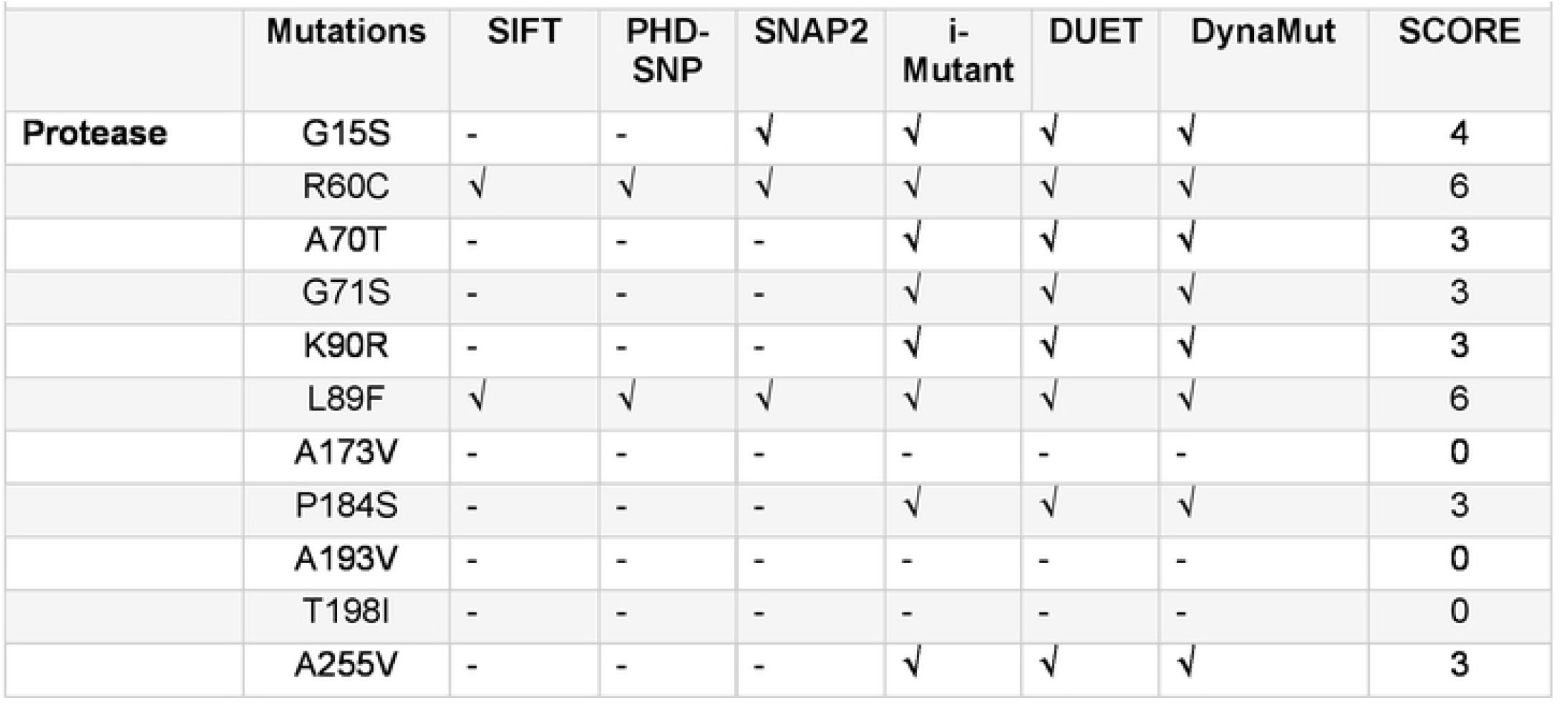
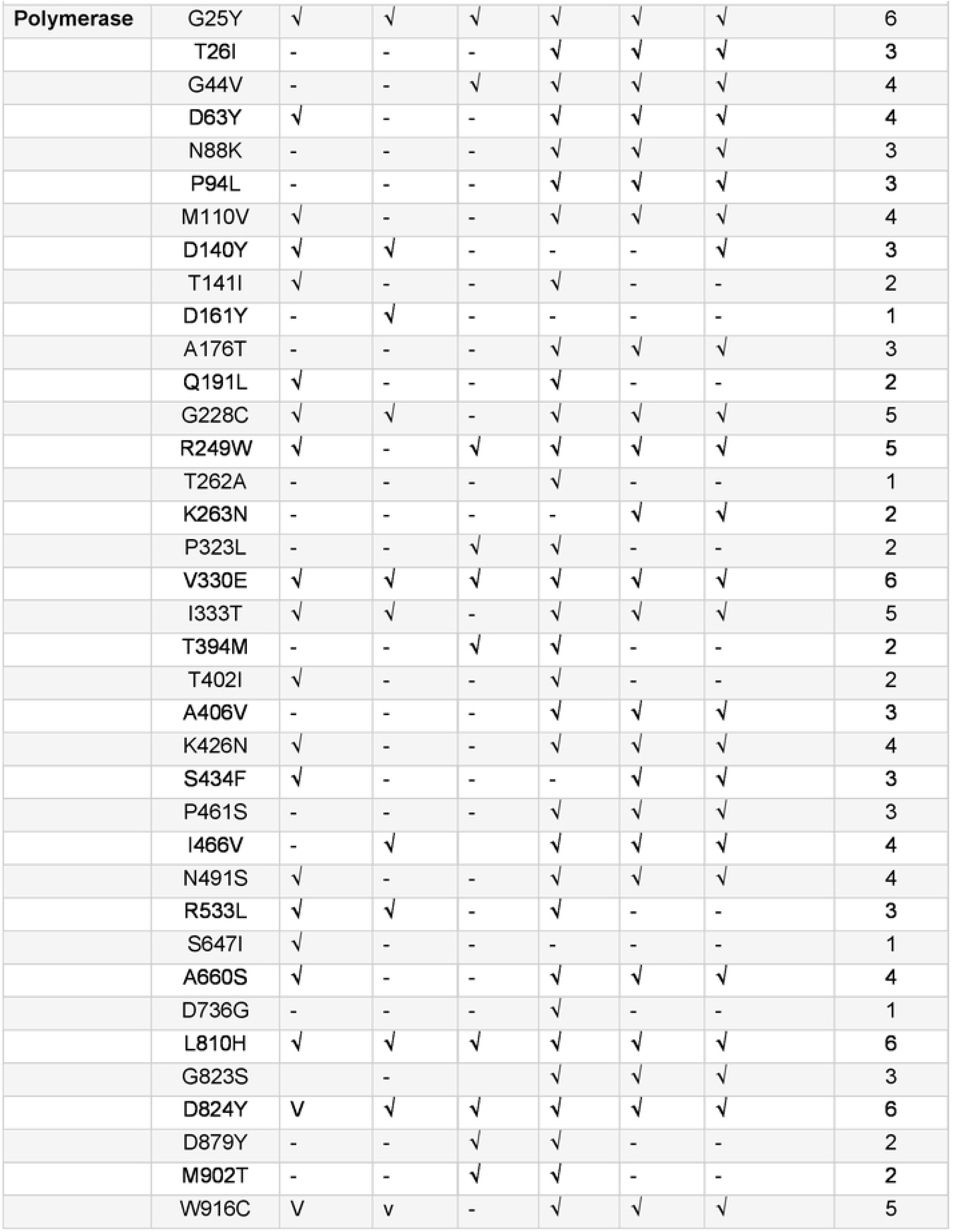
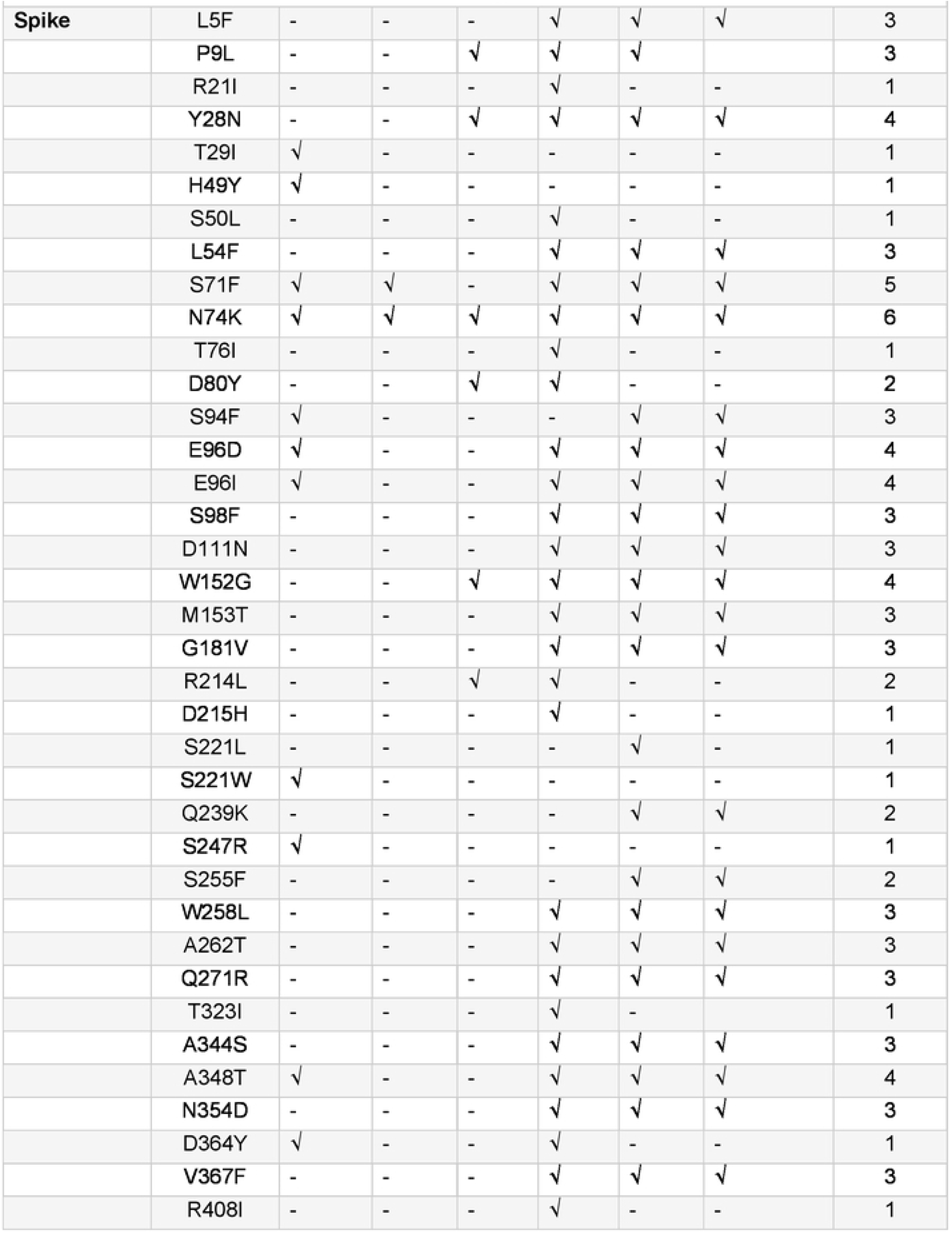
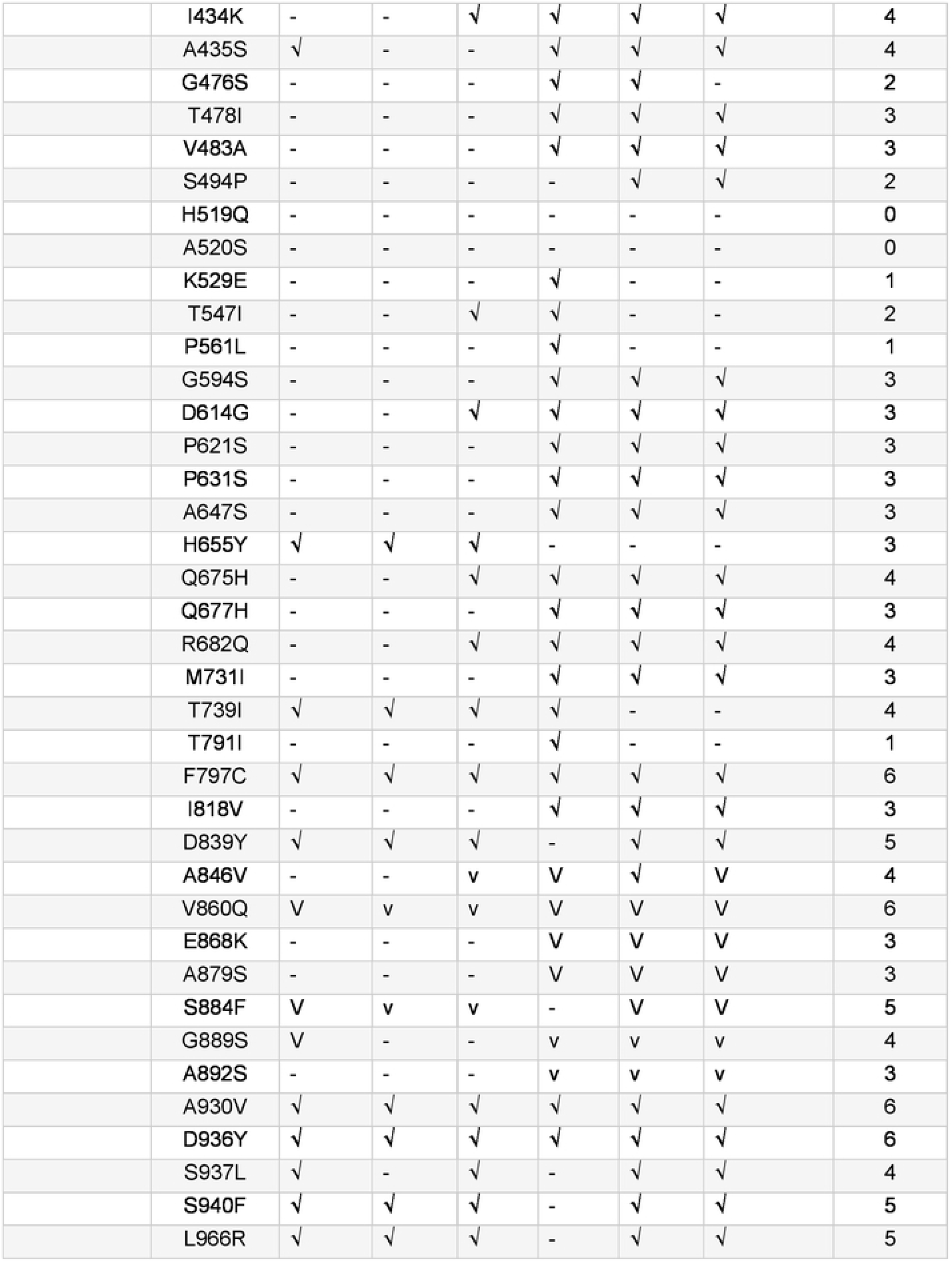
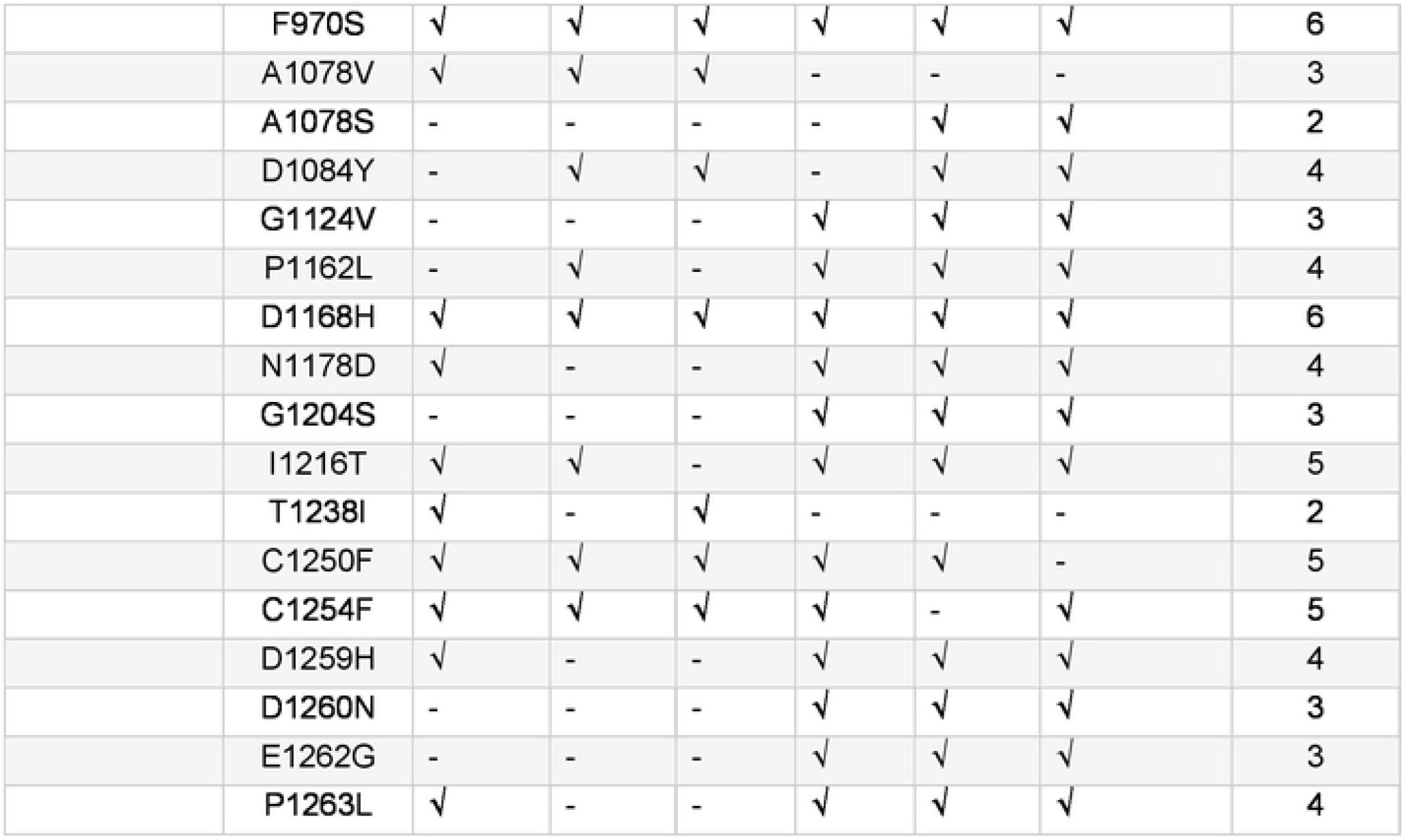
Prediction of deleterious mutations. Variations in 3CLpro **(A)**, RdRp **(B)**, and S **(C)** of SARS-CoV-2 that were predicted to be “deleterious” by all the six pieces of software.

**Figure 1.**
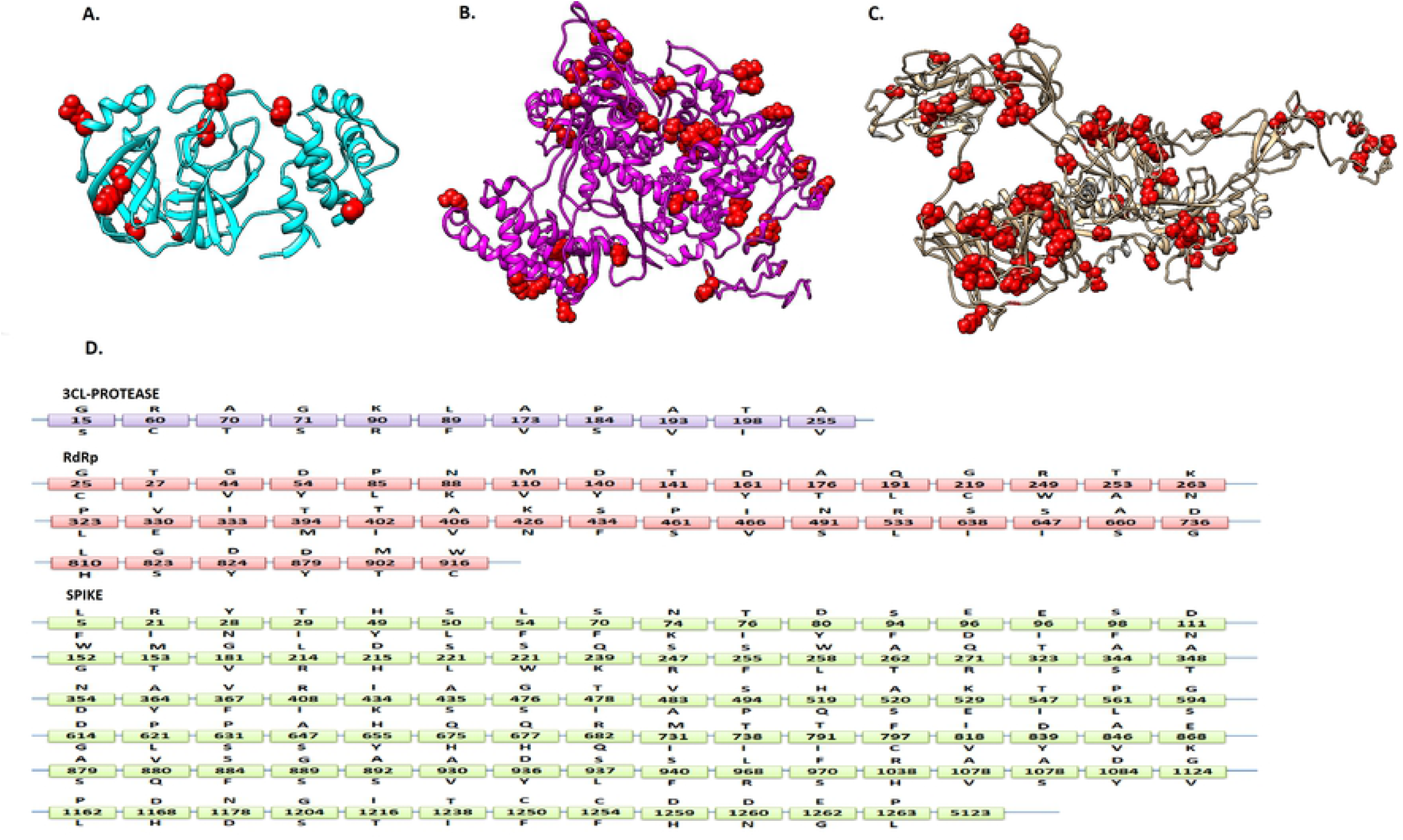
Mutation representation. Locations of 3CLpro **(A)**, RdRp **(B)**, and S **(C)** of SARS-CoV-2 mutations are presented in red spheres. **(D)** The letter(s) above the box refer the wild type amino acid and the letter(s) below the box are relevant substitutions reported in this study.

To further evaluate the effect of the given mutations on the structure and function of respective proteins, a variety of *in-silico* SNP prediction algorithms were used. NC_045512 was taken as a wild type genome. Its S and RdRp structure were predicted by i-TASSER, whereas crystal structure of SARS-CoV-2 3CLpro was retrieved from PDB (PDB ID: 5re5).

### Analyzing the effect of mutations on structural and functional stability of the proteins

Six prediction software tools, including SIFT [10], PhDSNP [15], SNAP2 [12], I-Mutant version 2.0 [11], DUET [14], and Dynamut [13] were employed to predict the effects of total 140 mutations in S (92), RdRp (37), and 3CLpro (11). According to SIFT analysis, in the S protein, 34 mutations were found to be deleterious and 58 mutations tend to be tolerated (neutral) in nature. In the RdRp protein, 20 mutations were declared in-tolerated, while 17 were tolerated. In the 3CLpro protein, two mutations were predicted as in-tolerated and nine mutations were tolerated.

PhD-SNP predicted 20 mutations in S protein as damaging or deleterious, 11 in RdRp, and two in 3CLpro protein. SNAP2 revealed that 29 mutations in S, 10 in RdRp, and three in 3CLpro could affect the overall function of these viral proteins. It also predicts which type of amino acid that affect the function of the protein when altered at a particular position. Based on its predicted analysis, a heat map is also generated depicting the abilities of the amino acids to change the function of respective viral proteins (Figure 2A).

**Figure 2.**
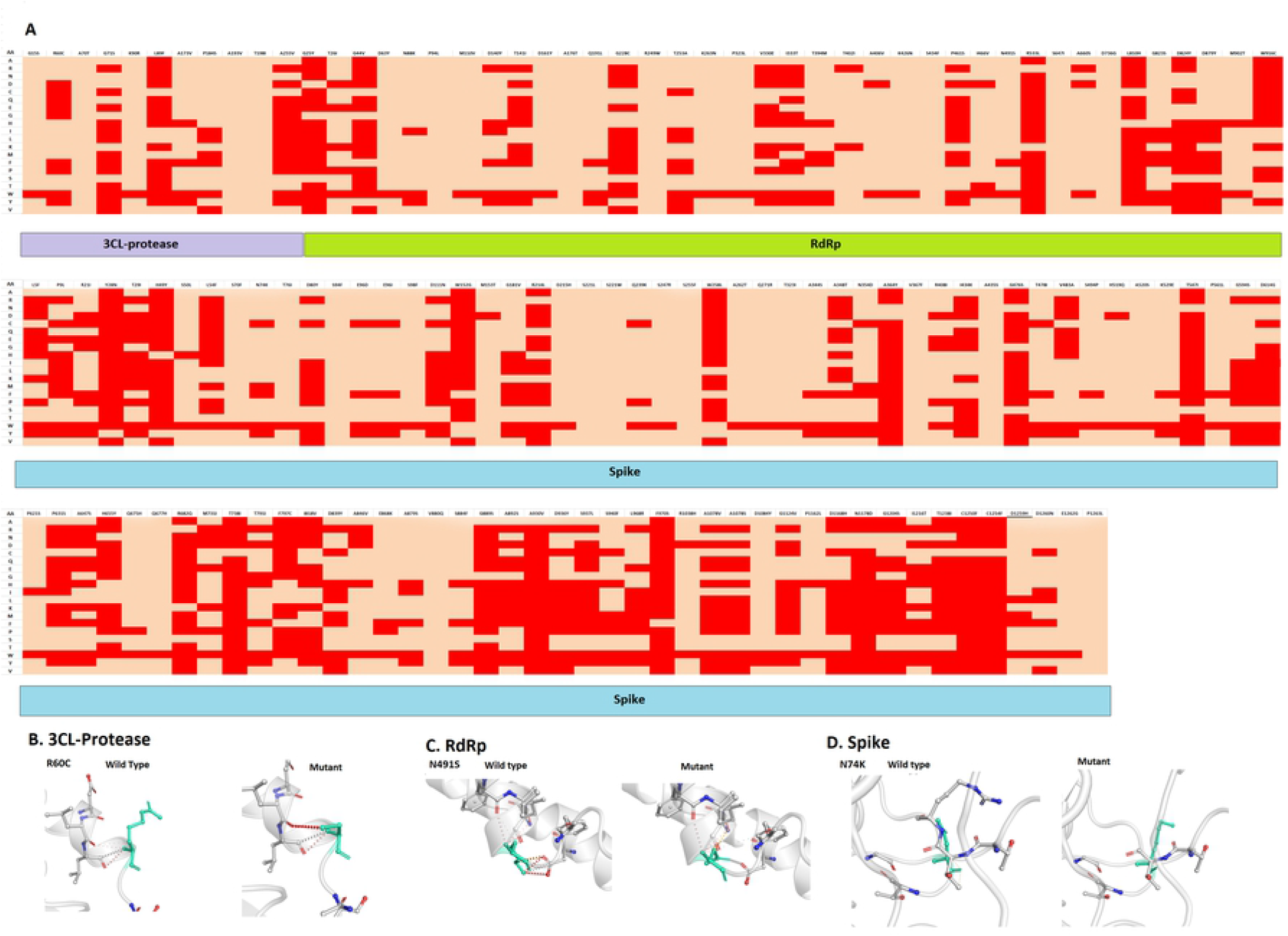
Heatmap representation of deleterious and non-deleterious mutations. **(A)** Heatmap representation showing possible substitution at each mutation position. Dark red indicates a high score (strong signal for effect), and green showed a low score (strong signal for neutral/no effect) based on SNAP2 analysis. **(B-D)** The effects of mutations (R60C, 3CLpro; N491S, RdRp; and N74K, S) on the structural stability of viral proteins predicted by Dynamut web server.

Findings of i-Mutant showed that out of 92 mutations, 71 were deleterious for the S structure. It also revealed that 32 mutations in RdRp and seven in 3CLpro are deleterious mutations. According to DUET, 68 mutations in S, 23 mutations in RdRp, and eight mutations in 3CLpro proteins are deleterious in nature. Findings of Dynamut suggest that 65 mutations in S, 25 in RdRp, and eight in 3CLpro can affect the structural conformation of the respective viral proteins. It also predicts interatomic interactions of wild-type and mutant amino acid with the environment based on atom type, interatomic distance, and angle constraints. Some of the selected deleterious mutations of S, RdRp, and 3CLpro as well as interatomic interaction analysis have been shown in Figure 2B-2D.

Details of all predicted mutations and their possible effects on the encoded proteins have been demonstrated in Table 1. These analyses predict mutations that could affect the structural stability of protein by changing the flexibility and rigidity in the targeted proteins. To evaluate these mutations, six tools have been employed, each tool has different strategies and parameters to predict deleterious mutations. The mutations with more positive results were more likely to be truly deleterious. Mutations observed to be deleterious by more than three prediction algorithms have been classified as high-risk (see Material and Methods).

Figure 3 shows the prediction results of six computation tools. As a result, five mutations were predicted to be neutral with a score of 0, while 19, 17, 49, 25, 12, and 13 mutations got a score of 1, 2, 3, 4, 5 and 6, respectively (Figure 3). Based on the given criteria, 32 mutations in S, 15 in RdRp, and three in 3CLpro (Table 1) met these criteria (score ≥ 4) and were chosen for further analysis (Figure 3). Among these pathogenic mutations, D614G (score=4) in S region has already been reported to be associated with a greater infectivity [16]. Another highly prevalent mutation (P323L) in RdRp region was found to be neutral (score=2), whereas its infectivity has not been reported so far. Finally, all these deleterious mutations were mapped on 3D structure of the viral proteins. It was observed that all these mutations were uniformly distributed on the viral protein structures.

**Figure 3.**
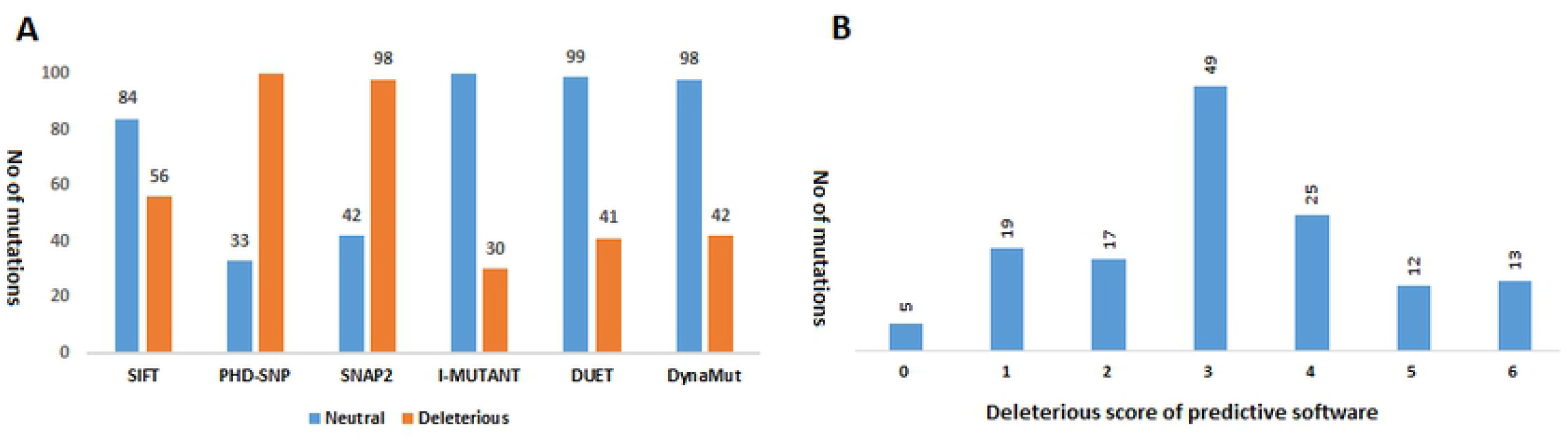
Prediction of pathogenicity of nsSNPs by SIFT, PhD-SNP, SNAP2.0, I-MUTANT, DUET, and DynaMut software. **(A)** The number of “deleterious” or “neutral” protein variants predicted by each bioinformatics tool. **(B)** Number of protein variants with different scores of six bioinformatics tools.

### Localization of the deleterious mutations within the binding sites of viral proteins

The 3D structure SARS-CoV-2 protease was retrieved from PDB with PDB ID 5RE5. For S and RdRp proteins, top i-TASSER predicted models were selected on the basis of C-score. RAMPAGE and ProSA web servers were further used to verify the reliability of predicted models.

The results of the predicted 3D RdRp model showed 83% of residues in favored region, 10.8% in additional allowed region, and 6.2% in outlier region. Tertiary structure of S protein showed 75.2% in favored region, 14.8% in allowed regions, and 10% in outlier regions that highly indicates a good stereo-chemical quality of the predicted structures. By using these 3D structures, COACH and CASTP servers predicted the possible ligand binding sites of these proteins. Ligand binding sites predicted by both servers were considered as potential binding sites. It was observed that in S protein, 22 out of 37 deleterious mutation positions including 28, 71, 74, 96, 152, 348, 435, 675, 682, 797, 824, 846, 860, 930, 936, 970, 1168, 1178, 1168, 1250, 1258, and 1259 were lying in the ligand binding site. In RdRp, 13 predicted deleterious mutation positions (25, 44, 63, 110, 228, 249, 333, 426, 491, 660, 810, 824, and 916) were lying in the ligand binding sites. While in 3CLpro, all selected deleterious mutation positions (15, 60, and 89) lie within the binding site.

### Normal mode analysis of highly deleterious mutations

iMODs is a user-friendly interface for normal mode analysis methodology. It provides the detailed information about mobility (B-factors), eigenvalues, covariance map, and deformability. The eigen value represents the total mean square fluctuations and is related to the energy required to deform the structure. The lower eigenvalues represent the easier deformation of the protein. iMODs analysis revealed that all selected deleterious mutations decrease the eigen values of RdRp, S, and 3CLpro proteins, indicating the deleterious effects of the evolving mutations in the selected viral proteins (Figure S1).

## Discussion

The current study is based on *in-silico* mutagenesis analysis of SARS-CoV-2 RdRp, S, and 3CLpro proteins to identify mutations and their possible structural and functional impact to the encoded viral proteins. In this study, 92 mutations in S, 37 in RdRp, and 11 in 3CLpro proteins have been identified in the available sequence data from all over the world. The effect of such mutations on the structure and function of respective viral proteins is important to predict the evolutionary potential of the viral proteins. However, *in-silico* prediction of the impact of aminoacid variants to the proteins’ structure and function may, sometimes, be considered as an alternative or pre-study indicator to *in vitro* expression level studies [17]. In addition, interpretation of the proteomic variants in the light of their phenotypic effects is one of the emerging crucial tasks to solve in order to advance our understanding of how these variants affect SARS-CoV-2 proteins structural and functional behavior. RdRp, S, and 3CLpro proteins of SARSCoV-2 are important targets for antiviral drug and vaccine development [18] and thus, have been selected for bioinformatics analysis in this study. Any mutation in these viral proteins could be either beneficial or deleterious for the virus in this pandemic [3]. Therefore, we identified mutations in the selected viral proteins as well as the possible impact of these mutations on the overall structure and function of these proteins.

It was observed that most of the mutations were lying in the S region (92), followed by RdRp (37), and 3CLpro (11). Highly mutated amino acid was observed at the position of D614G (50%) in S protein and P323L (49%) in RdRp protein. By using various *in silico* algorithms and selecting scoring criteria (0-6), it was estimated that 32 mutations in S, 15 in RdRp, and 3 in 3CLpro proteins were deleterious in nature and probably affect the overall structure and function of these viral proteins. Among these mutations, D614G is highly prevalent and associated with greater infectivity of COVID-19 infection. It was also found to be pathogenic in nature (score=4), thus validating the current results. Another highly prevalent mutation P323L in RdRp was found to be neutral (score=2). Similarly, the remaining mutations are rare and does not appear to be more deleterious.

Together, these findings have implications for our understanding of SARS-CoV-2 mutations. These mutations do not only affect structural and functional abilities of viral proteins, but they might also affect the binding affinities of these viral proteins with various drugs, as most of these pathogenic mutations are also present in ligand binding regions. This characterization of drug and vaccine target protein variants of SARS-CoV-2 could help us in understanding the pathogenesis, treatment options, vaccines design, and diagnostic strategies. It would potentially be significant to characterize the impact of these identified pathogenic mutations by employing various *in vitro* and molecular approaches.

## Author Contributions

**Conceptualization:** Aqsa Ikram

**Data curation:** Aqsa Ikram

**Formal analysis:** Aqsa Ikram, Anam Naz, Faryal Mehwish Awan

**Investigation:** Aqsa Ikram, Anam Naz, Faryal Mehwish Awan, Bisma Rauff, Ayesha Obaid

**Methodology:** Aqsa Ikram, Arif Malik, Mohamad S. Hakim

**Project administration:** Bisma Rauff, Ayesha Obaid

**Supervision:** Arif Malik, Mohamad S. Hakim

**Writing – original draft:** Aqsa Ikram

**Writing – review & editing:** Aqsa Ikram, Mohamad S. Hakim.

## Funding

This study received no external funding.

## Acknowledgments

We are grateful to the Institute of Molecular Biology and Biotechnology (IMBB), The University of Lahore (UOL), Lahore, Pakistan for their administrative support.

## Declaration of Competing of interest

The author declares no conflict of interest.

## Figure Legends

**Supplementary Figure 1**. Normal mode analysis of WT (**A**) and mutant (L89F) (**B**) 3CLPRO protein. Detailed profiles of mobility (B-factors), eigenvalues and deformability have been shown.

